# *De novo* expression of neuropeptide Y in sensory neurons does not contribute to peripheral neuropathic pain

**DOI:** 10.1101/2025.01.25.634591

**Authors:** A.H. Cooper, A. Nie, N.S. Hedden, H. Herzog, B.K. Taylor

**Affiliations:** Pittsburgh Center for Pain Research and Opioid Research Center, Department of Anesthesiology & Perioperative Medicine, Pittsburgh, PA, 15213, USA; St Vincent’s Centre for Applied Medical Research (AMR), Sydney; St Vincent’s Clinical School, Faculty of Medicine, The University of New South Wales, Sydney, Australia

**Keywords:** Allodynia, hyperalgesia, nerve injury, regeneration, BIBO3304, gabapentin

## Abstract

Nerve damage induces a robust *de novo* expression of the pain-modulatory peptide neuropeptide Y (NPY) in large-diameter primary afferent neurons that innervate the dorsal horn of the spinal cord and the dorsal column nuclei. To determine whether this functions to modulate peripheral neuropathic pain, we selectively deleted the *Npy* gene in neurons of the dorsal root ganglion (DRG), without disruption of its expression in brain or dorsal horn neurons. We then subjected sensory neuron-specific NPY deletion mutant mice (*Pirt*-NPY) and their wild-type controls to either sham surgery, spared sural nerve injury (SNI) or spared tibial nerve injury (tSNI). Conditional *Npy* deletion did not change the severity or duration of static mechanical, dynamic mechanical, or cold allodynia in SNI or tSNI models, nor ongoing neuropathic pain as assessed with conditioned place preference to gabapentin. When injected after the resolution of tSNI-induced mechanical hypersensitivity (a latent pain sensitization model of chronic neuropathic pain), the NPY Y1 receptor-specific antagonist BIBO3304 equally reinstated mechanical hypersensitivity in *Pirt*-NPY mice and their wildtype controls. We conclude that nerve injury-induced upregulation of NPY in sensory neurons does not cause mechanical or cold hypersensitivity or ongoing pain, and that tonic inhibitory control of neuropathic pain by NPY in the spinal cord is mediated by release from dorsal horn interneurons rather than sensory neurons.

## INTRODUCTION

Neuropeptide Y (NPY), first described in 1982 as a 36 amino acid peptide with five tyrosine residues in its primary structure^1^, regulates myriad physiological processes including feeding, cardiovascular function, and emotions throughout the nervous systems of rodents and humans. NPY is a principally inhibitory signaling molecule, as all NPY receptor subtypes (Y1,Y2,Y4, and Y5) are G_i_PCRs. Preclinical research from the past three decades has revealed that NPY plays a major role in pain modulation by acting at Y1 and Y2 receptors throughout the central and peripheral nervous systems.

Within the peripheral somatosensory nervous system, primary afferent neurons (PANs) of the dorsal root ganglia (DRG) express minimal numbers of NPY transcripts or levels of peptide under normal conditions. After peripheral nerve injury, however, *Npy* is dramatically up-regulated relative to other genes^2^. This is particularly evident in the somata of medium- and large-sized PANs^3,4^ as well as their central terminals within deeper laminae III-V of the dorsal horn ^5^ and dorsal column nuclei (DCN)^6^. However, the functional contribution of *de novo* NPY in the modulation of pain is unclear, largely because NPY expressed by primary afferents can be released at sites at which NPY is either pronociceptive ^7-9^ or antinociceptive^9-11^. Pronociceptive effects of NPY are observed in the periphery, DRG, and the dorsal column nuclei (DCN). For example, injection of NPY into the nucleus gracilis (the DCN innervated by hindpaw afferents)^8^, DRG ^9^ or hindpaw^12^ induced behavioral signs of mechanical hypersensitivity. Administration of Y1 or Y2 antagonists into the DRG and nucleus gracilis attenuated behavioral signs of neuropathic pain. Together, these data indicate a pronociceptive role of exogenous and endogenous NPY in these regions.

Conversely, NPY in the spinal cord dorsal horn and various brain regions (such as the periaqueductal grey, parabrachial nucleus, and rostroventral medulla) is a powerful inhibitor of hyperalgesia and allodynia in rodent models of inflammatory and neuropathic pain^11,13,14^. In the dorsal horn, Y1 is primarily expressed by excitatory interneurons whereas Y2 is largely restricted to the terminals of a subpopulation of afferents^15-17^. Intrathecal injection of NPY, though not analgesic in the naïve state, is antihyperalgesic following SNI^17-19^. Both Y1 and Y2 antagonists block this effect^18^. Furthermore, intrathecal administration of Y1 or Y2 antagonists reinstated neuropathic pain-like behaviors in the tSNI variant of the SNI model^11^. These results indicate antihyperalgesic effects of exogenous NPY and long-lasting endogenous NPY release in the spinal cord, the latter of which could originate from: 1) intrinsic terminals of inhibitory NPY interneurons in the superficial dorsal horn; and/or 2) volume transmission from the central terminals of damaged primary afferent neurons. Volume diffusion could explain how NPY release from deeper layers reach densely packed Y1R-expressing interneurons in laminae I-II ^15,17^. As these neurons are necessary and sufficient for the expression of inflammatory and neuropathic pain ^14,17,20,21^, it is plausible that NPY released by primary afferent neurons could bind to Gi-coupled receptors on Y1-expressing neurons and thereby decreased behavioral signs of chronic pain.

The functional consequences of nerve injury-induced up-regulation of NPY in primary afferent neurons, notably in the modulation of neuropathic pain, has been a long-standing question^22^ that has proven difficult to answer in the absence of a rigorous approach to selectively eliminate NPY from PANs while leaving NPY-expressing spinal cord interneurons intact. To address this gap, we generated transgenic mice with loxP sites flanking the Npy gene (*Npy*^lox/lox^), crossed them with *Pirt*^Cre^ mice to selectively knockout NPY from primary afferent neurons, and then evaluated behavioral signs of mechanical and cold hypersensitivity and spontaneous pain in these *Pirt*-NPY mice and their wild-type controls after sham or nerve injury.

## MATERIALS AND METHODS

### Animals

All procedures were approved by the University of Pittsburgh Institutional Animal Care and Use Committee in accordance with American Veterinary Medical Association and International Association for the Study of Pain guidelines. Male and female mice aged 7-16 weeks at the beginning of experiments were housed 2-4 per cage and maintained on a 12/12 light/dark cycle at 20-22°C and 45 ± 10% relative humidity, with food and water provided *ad libitum*. Mice were handled and habituated to testing equipment for 30 min/day for 3 consecutive days prior to experimental manipulations and all procedures were performed during the animals’ light cycle (between 7am and 7pm).

NPY floxed (NPY^lox/lox^) mice generated on a pure C57Bl/6 background were a gift from Prof Herbert Herzog (Garvan Institute of Medical Research, Australia)^23,24^. Pirt^cre^ mice (cre recombinase expression driven by the *Pirt* promotor, expressed in ∼95% of primary afferent neurons) were a gift from Dr Xinzhong Dong (Johns Hopkins University, USA)^25^. Both lines were maintained at the University of Pittsburgh. Mice with primary afferent neuron-specific knockout of NPY (Pirt^cre^::NPY^lox/lox^) were generated by crossing NPY^lox/lox^ with Pirt^cre^ mice; the breeding strategy is illustrated in **Fig 1a**. Homozygous NPY^lox/lox^ or Pirt^cre^::NPY^lox/+^ littermates were used as controls. Successful deletion of NPY expression was confirmed by PCR and immunohistochemistry.

**Figure 1.**
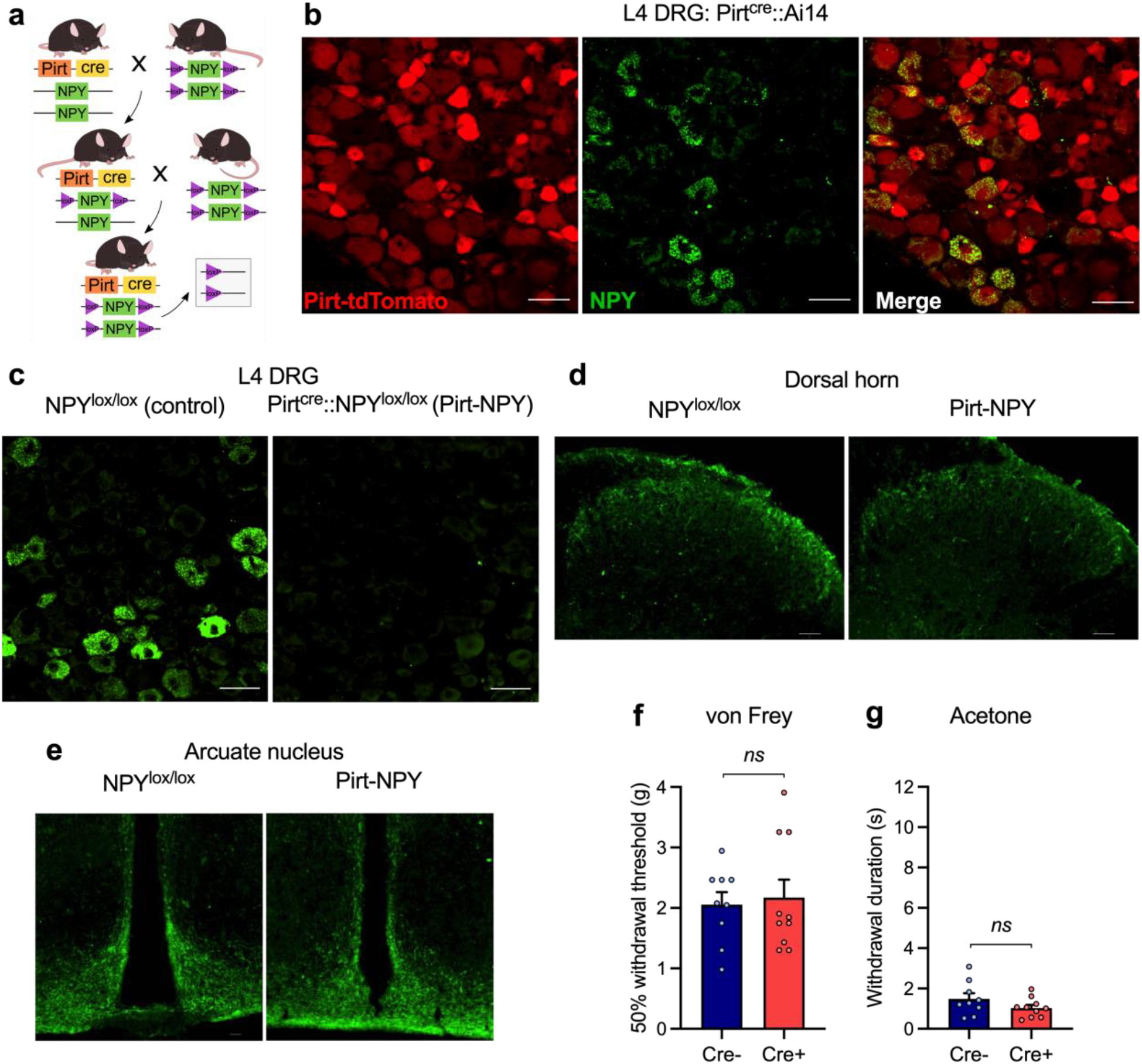
*Pirt*-NPY mice do not express nerve injury-induced up-regulation of NPY in DRG neurons. **(a)** Schematical illustration representing the breeding strategy for primary afferent neuron-specific knockout of NPY: Pirt^Cre^::NPY^lox/lox^ (NPY cKO) and the knockout allele generated in the presence of the cre transgene. **(b)** Representative image of L4 DRG neurons from a Pirt^cre^::Ai14 mouse 14 days following SNI, showing that NPY is expressed by Pirt-positive neurons. Scale bars = 50 μm. **(c)** Representative image showing NPY expression in L4 DRG neurons 14 days after tibial spared nerve injury (tSNI) in NPY^lox/lox^ controls (left) and *Pirt*-NPY mice (right). Scale bars = 50 μm. **(d and e)** Representative images showing dorsal horn **(d)** and arcuate nucleus **(e)** NPY expression is conserved in the CNS of *Pirt*-NPY mice. Scale bars = 50 μm. **(f and g)** In the absence of injury, Pirt^Cre^ expression did not alter behavioral hindpaw responses after **(f)** mechanical or **(g)** cold stimulation. Paired t-tests; mechanical: t_17_ = 0.31, *P* = 0.76; cold: t_17_ = 1.45, *P* = 0.16; *n* = 9-10). Data represents mean ± SEM.

### Sural and tibial spared nerve injury surgeries

Sural (SNI) and tibial (tSNI) spared nerve injury surgeries were performed as previously described ^11,26^. Anesthesia was induced with 5% isoflurane (Abbott Laboratories, USA) and then maintained at 2%. The left hindpaw was secured with tape in hip abduction and the operative field (lateral surface of the thigh) was shaved. Ophthalmic ointment was applied to the eyes and the shaved area was swabbed with chlorhexidine solution (Chloraprep, BD Healthcare, USA). A longitudinal incision was made in the skin at the lateral mid-thigh. Using blunt dissection an opening was made through the biceps femoris, exposing the sciatic nerve and the three peripheral branches (sural, tibial and common peroneal nerves). Using 6-0 silk suture (Ethicon, USA), the common peroneal and tibial nerves (SNI), or common peroneal and sural nerves (tSNI) were ligated, cut 1-2 mm on either side of the suture using spring scissors and a small piece of each nerve was removed. The muscle was closed with two 5-0 Vicryl sutures (Eithcon, USA), the skin incision closed with two 10 mm wound clips (Alzet, USA) and topical Neosporin ointment (Johnson and Johnson, USA) was applied to the wound. Sham-operated mice received isoflurane for the same duration as SNI mice but had no incision. Wound clips were removed ten days after surgery.

### Drug administration

#### Intrathecal injection

Mice were shaved over the lumbar spine (∼20 × 20 mm) and acclimated to restraint to minimize stress during intrathecal injection. Awake mice were gently restrained using a towel and secured at the pelvic girdle with experimenter thumb and forefinger. The Y1R-specific antagonist BIBO3304 (Tocris, UK) was dissolved in saline, and injections were administered at a volume of 5 µL, between L4 and L5 vertebrae using a 30-G needle attached to a 25 µL Hamilton syringe ^27^. Successful entry of the needle into the intradural space was indicated by observing a tail flick. The needle was then secured while the drug was slowly injected over 10 seconds and held in place for an additional 10 seconds to minimize backflow upon removal of the needle.

#### Intraperitoneal injection

For conditioned place preference experiments, gabapentin (100 mg/kg; Sigma-Aldrich, USA; dissolved in saline) or vehicle was administered by intraperitoneal injection at a volume of 10 mL/kg. Mice were restrained by scruff and the drug was injected into the lower quadrant of the abdomen using a 27-G needle.

### Behavioral testing

#### Von Frey assessment of static mechanical allodynia

Hindpaw 50% mechanical withdrawal thresholds were measured using the von Frey (vF; Stoelting, Inc, IL, USA) up-down method ^28^. Mice were placed in an acrylic box, opaque on all sides, atop an elevated wire mesh platform, for at least 15 min prior to assessment. The stimulus was applied to the hindpaw at the lateral aspect of the glabrous skin, for mice that received SNI surgery, or at the center of the glabrous skin between the tori, for tSNI or sham mice. Each paw was stimulated using a predefined set of 8 monofilaments (0.008 to 6 g), starting each trial with an intermediate filament (0.16 g), applied perpendicular to the skin, causing a slight bending, for 3 s. In case of a positive response (rapid withdrawal or licking of the paw within 3 s of removing of the filament, but ignoring normal ambulation or rearing), the next smallest filament was tested, with at least 2 min between stimulations. In case of a negative response, the next larger filament was tested. Each trial continued until 4 measurements beyond the first change in response (i.e., no response then response, or vice versa) were taken. 50% mechanical withdrawal thresholds were calculated using the statistical method described by ^29^.

#### Cotton swab assay of dynamic mechanical allodynia

Sensitivity to a dynamic light touch stimulus was assessed using a cotton swab as described previously^30,31^. Mice were acclimated for at least 30 min to the same acrylic boxes and elevated wire mesh platform as for von Frey testing. The tip of a cotton swab (Fisherbrand wood/cotton swab; Fisher Scientific, USA) was teased apart to approximately three times the original size. Using a sweeping motion, the swab was brushed gently across the plantar surface of the left hindpaw in a medial-to-lateral direction. Care was taken to apply the same force (3 ± 0.5 g when applied to a scale) and velocity (sweeping motion completed so swab is in contact with hindpaw for ∼0.5 s) during all stimulations. Scoring: 0, no reaction to the swab; 1, a rapid, single withdrawal; and 2, a withdrawal accompanied by paw fluttering for >0.5 s and/or licking or biting of the stimulated paw. Five applications were performed per timepoint, with at least 2 min between each, and the mean response score for each mouse was calculated.

#### Acetone assay of cold allodynia

To test sensitivity to a cooling stimulus, the acetone assay was used as described previously. A piece of PE-90 tubing, melted at the distal tip and attached to a 5 mL syringe, was used to apply a drop (10–12 ul) of acetone to the lateral aspect (SNI) or center (tSNI or sham) of the hindpaw skin, ipsilateral to injury. The duration of time that the mouse spent raising or licking this paw was recorded, with a cut-off of 30 sec. Three measurements were averaged per time point, with at least 5 min between acetone applications.

#### Conditioned place preference

To test for the affective / spontaneous component of pain, a three-day conditioning protocol used a biased chamber assignment to test for conditioned place preference (CPP), as described previously ^32^. On the acclimation day (Day 1,seven weeks after SNI or sham surgery), mice had free access to explore all chambers of a 3-chamber conditioned place testing apparatus (2 conditioning side chambers: 170 × 150 mm and a center chamber: 70 × 150 mm; height: 200 mm; San Diego Instruments, USA) for 30 minutes. Mice were able to discriminate between chambers using visual (vertical versus horizontal black-and-white striped walls) and sensory (rough versus smooth textured floor) cues. For pre-conditioning (Days 2 and 3), mice were again allowed to freely explore for 15 min whilst their position was recorded via a 4 × 16 infrared photobeam array by associated software (San Diego Instruments, USA). To avoid preexisting chamber bias, mice spending more than 80% (720 s) or less than 20% (180 s) of time in either side chamber during preconditioning were excluded from further testing. For conditioning (days 4-6), each mouse’s preferred chamber was paired with saline and non-preferred chamber with gabapentin (biased design). Each morning, mice received a i.p. saline injection, were returned to their home cage for 5 min, then placed in the designated side chamber for 30 min. 4 hours later, mice received i.p. gabapentin (100 mg/kg; Sigma-Aldrich, USA; dissolved in saline), returned to their home cage for 5 min, then placed in the gabapentin-paired chamber for 30 min. On test day (Day 7), mice could freely explore whilst their position was recorded, as during pre-conditioning, for 15 min. Difference scores were calculated as the time spent in each chamber on test day minus time spent during pre-conditioning. The total time spent in each conditioning chamber during pre- and post-conditioning were also compared for analysis.

### Histology

#### Immunohistochemistry

Mice were anesthetized with an overdose of pentobarbital (200 mg/kg) and transcardially perfused with 4% paraformaldehyde (PFA). Brains, spinal cords and lumbar segments 3 to 5 dorsal root ganglia ipsilateral to injury were removed and post-fixed in 4% PFA overnight, cryoprotected in 30% sucrose for a further 48 hours and then embedded in optimal cutting temperature media (OCT; Tissue Tek, Andwin Scientific, USA). Using a cryostat (Cryostar NX70, Fisher Scientific, USA), 10 µm sections of DRG were collected on-slide (Superfrost+, Fisher Scientific, USA), and 30 µm sections of spinal cord or brain sections were collected free-floating in 0.1 M PBS. Sections were washed in PBS, then blocked in PBS containing 3% normal goat serum (NGS; MP Biomedicals) and 0.3% Triton X-100 (VWR, USA) for 1 hour, and incubated for 18 hours at room temperature with rabbit anti-NPY antibody (1:1000; Peninsula Laboratories, CH; Cat# T-4070, RRID: AB_518504), diluted in 1% NGS and 0.3% Triton X-100. Following washes, sections were incubated with Alexa Fluor-488-conjugated goat anti-rabbit IgG (H+L) (1:1000; Invitrogen, USA; Cat# A11011; RRID: AB_143157) for 1.5 hours. After further washes, brain and spinal cord sections were mounted on gelatinized slides, and all slides were rinsed briefly in distilled water, dried, and coverslipped with Vectashield antifade mounting medium with DAPI (Vector Labs, USA; Cat# H-1500-10).

#### Imaging

Sections were imaged with a Nikon Ti2 inverted epifluorescence microscope equipped with 10x (0.45 NA) and 20x (0.75 NA) objectives, slide scanner and a Prime BSI camera (Photometrics, USA), controlled using NIS Elements (Nikon, Japan). The same exposure time (100 to 300 ms) was used for all images captured in each channel. Two to nine images encompassing the field of view were captured and stitched online with 10% overlap. Brightness and contrast were adjusted in the same manner for each set of images of a structure (DRG, dorsal horn, or arcuate nucleus).

### Statistical analyses

All behavioral and immunohistochemical results are presented as mean ± SEM. Statistical analyses were performed in Prism 9.2 (GraphPad Software Inc., USA) using Student’s (unpaired) t-test, 2-way or 3-way RM ANOVAs. If a main ANOVA effect was significant, then Tukey’s post-hoc tests were conducted. Separate 3-way RM ANOVAs examining allodynia over time were performed for SNI and tSNI groups versus Sham. The threshold for statistical significance was set at *P* < 0.05. CPP heat maps were generated using a two-dimensional kernel density estimation based on the XY coordinates of mouse location for the duration of testing, performed using R 3.4.0.

## RESULTS

### Peripheral nerve injury-induced up-regulation of NPY is selectively abolished in DRG neurons in Pirt^cre^::NPY^lox/lox^ mice

We crossed Pirt^cre^ and NPY^lox/lox^ mice to generate a conditional knockout (cKO) mouse line in which the gene encoding for NPY was deleted from primary afferent neurons (**Fig. 1a**). To determine whether DRG neurons co-express Pirt^cre^ and NPY, we evaluated NPY immunoreactivity in Pirt^cre^::Ai14 mice after SNI. NPY+ and Pirt-tdTomato+ profiles overlapped completely, thus confirming that SNI increased NPY expression in sensory neurons (**Fig. 1b**). We next determined the efficacy of conditional knockout in Pirt^cre^::NPY^lox/lox^ mice (*Pirt*-NPY) to decrease this nerve injury-induced up-regulation of NPY. We found that NPY immunoreactivity in medium to large diameter neurons in L4 DRG was abolished in *Pirt*-NPY mice (**Fig. 1c**).

As expected, NPY immunoreactivity was partially reduced in the spinal dorsal horn (DH) of *Pirt*-NPY mice (**Fig. 1d**), likely due to loss of NPY in the central terminals of primary afferent neurons. NPY immunoreactivity in arcuate nucleus (**Fig. 1e**) was similar in *Pirt*-NPY and control mice. These data indicate that sensory neuron-specific NPY gene deletion did not extend to the brain and was localized to the soma and terminals of PANs. We also verified that Pirt^cre^ expression did not change baseline sensitivity to mechanical (**Fig. 1f**) or cold (**Fig. 1g**) stimuli in uninjured mice.

### Conditional deletion of NPY from primary afferent neurons does not change nerve injury-induced mechanical or cold hypersensitivity

Nerve injury dramatically increases the expression of NPY in primary afferent neurons. Nerve injury has also been proposed to increase the tonic or stimulus-evoked release of NPY at the peripheral terminals of PANs, as well as at their central terminals in the spinal cord dorsal horn and nucleus gracilis. To test the hypothesis that NPY in PANs provides tonic inhibition of neuropathic pain, we evaluated behavioral signs of mechanical and cold hypersensitivity in *Pirt*-NPY or control mice in two models of peripheral nerve injury: SNI to assess persistent, maximal hypersensitivity; and tSNI to examine transient, sub-maximal hypersensitivity. SNI induced mechanical hypersensitivity at the paw ipsilateral to injury (**Fig. 2a**) but not contralateral hindpaw (**Fig. 2b**). SNI induced dynamic mechanical allodynia (**Fig. 2c**) and cold hypersensitivity (**Fig. 2d**) at the ipsilateral hindpaw (**Table 1**; 3-way RM ANOVAs; Time x Injury interaction). NPY cKO changed neither mechanical hypersensitivity, mechanical allodynia, or cold hypersensitivity (**Table 1**; 3-way RM ANOVAs; Time x Injury x Genotype interaction).

**Figure 2.**
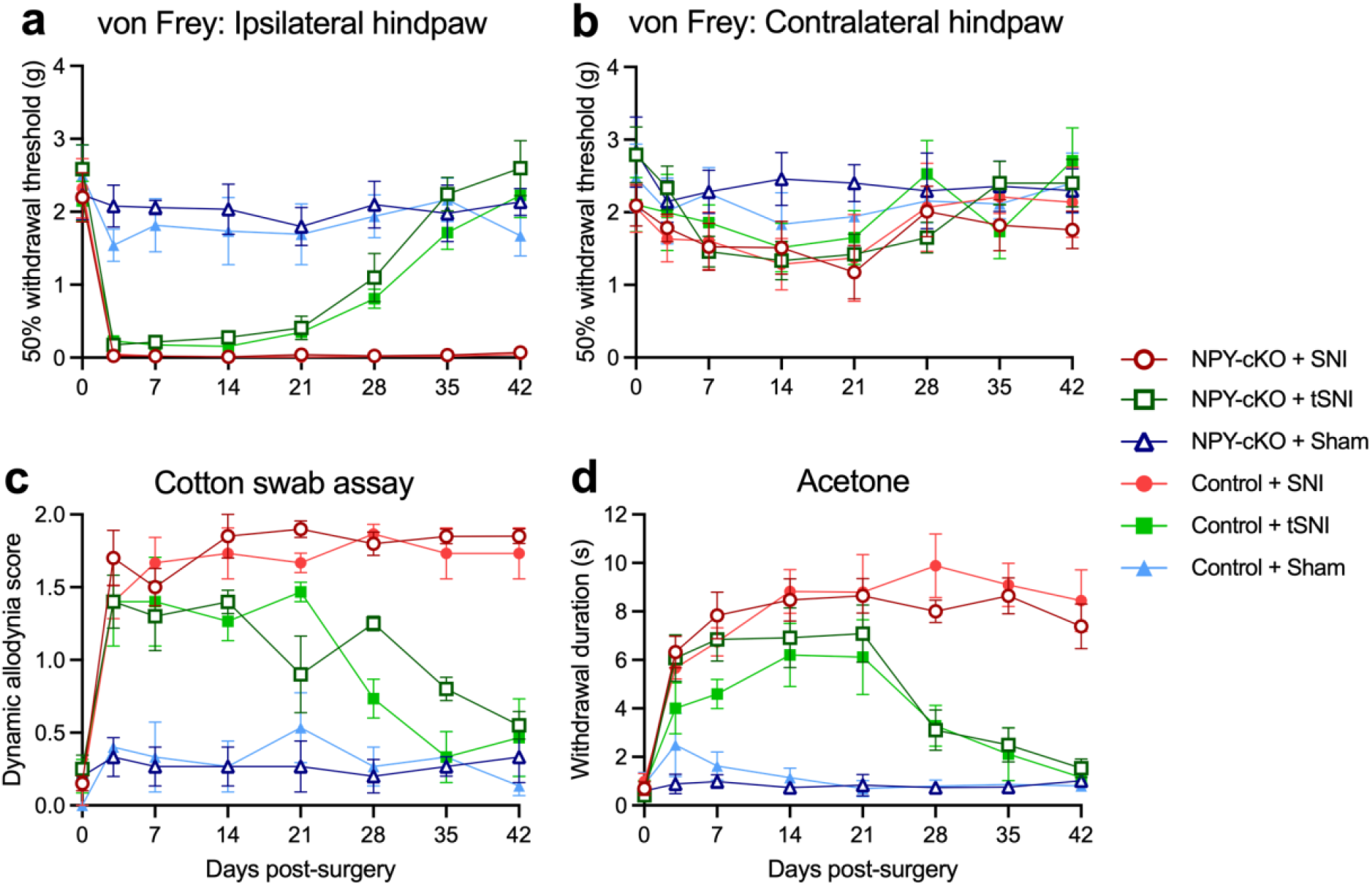
Sensory neuron-specific NPY cKO does not change mechanical or cold hypersensitivity after nerve injury. **(a** and **b**) Static mechanical hypersensitivity at the ipsilateral **(a)** and contralateral **(b)** hindpaws following SNI, tSNI or sham surgery in NPY^lox/lox^ controls and NPY cKO mice. **(c)** Dynamic mechanical allodynia at the ipsilateral hindpaw following SNI, tSNI or sham surgery in NPY^lox/lox^ controls and NPY cKO mice. **(d)** Cold sensitivity at the ipsilateral hindpaw following SNI, tSNI or sham surgery in NPY^lox/lox^ controls and NPY cKO mice. n=6-7. 3-way RM ANOVAs with Tukey’s post-tests: results in **Table 1**. Data represents mean ± SEM.

**Table 1.** 3-way RM ANOVA degrees of freedom and *P* values for analyses of Injury (sural or tibial spared nerve injury) and Genotype (control or NPY cKO) over time for static mechanical (von Frey; vF), dynamic mechanical (brush) or cold (acetone) allodynia. *n* = 6 – 7, except brush where *n* = 3 – 4.

|  | Sural spared nerve injury (SNI) |  | Tibial spared nerve injury (tSNI) |  |
| --- | --- | --- | --- | --- |
| Test | Time x Injury | Time x Injury x Genotype | Time x Injury | Time x Injury x Genotype |
| vF, ipsi | $F_{7,147} = 8.14, P < 0.001$ | $F_{7,147} = 0.36, P = 0.93$ | $F_{7,147} = 12.46, P < 0.001$ | $F_{7,147} = 0.81, P = 0.58$ |
| vF, contra | $F_{7,147} = 0.66, P = 0.70$ | $F_{7,147} = 0.14, P = 0.99$ | $F_{7,147} = 1.22, P = 0.30$ | $F_{7,147} = 0.77, P = 0.61$ |
| Brush | $F_{7,63} = 20.2, P < 0.001$ | $F_{7,63} = 1.11, P = 0.37$ | $F_{7,63} = 7.16, P < 0.001$ | $F_{7,63} = 1.18, P = 0.33$ |
| Acetone | $F_{7,147} = 25.4, P < 0.001$ | $F_{7,147} = 1.56, P = 0.15$ | $F_{7,147} = 14.5, P < 0.001$ | $F_{7,147} = 1.42, P = 0.20$ |

As with SNI, tSNI induced mechanical and cold hypersensitivity. As in previous studies^11^, this resolved within six weeks (**Fig. 2a-d**; **Table 1**). NPY cKO did not change the extent of mechanical or cold hypersensitivity (**Table 1**; 3-way RM ANOVAs; Time x Injury x Genotype interaction). Together, these results indicate that conditional deletion of NPY from PANs does not change the intensity of mechanical or cold allodynia following peripheral nerve injury.

### Conditional deletion of NPY from primary afferent neurons does not change the affective component of ongoing neuropathic pain

Clinical neuropathic pain includes not only hypersensitivity but also an ongoing pain associated with negative affective states ^33,34^. These states can be modelled in rodents with conditioned place assays ^21,32,35^. To test the hypothesis that primary afferent NPY tonically contributes to or attenuates affective pain after spared nerve injury, we evaluated conditioned place preference (CPP) to the pain relief provided by gabapentin (100 mg/kg, i.p.) according to the timeline illustrated in **Fig. 3a**. Prior to conditioning, mice displayed no preference to either side chamber (**Fig. 3b**; paired t-test; t_22_ = 0.98, *P* = 0.34; *n* = 23). Gabapentin induced CPP in mice that had undergone SNI but not sham surgery, indicating relief from the affective aspect of ongoing pain (**Fig. 3c-e**; 3-way ANOVAs with Tukey’s post-tests; Difference score Drug x Injury interaction: F_1,18_ = 21.33, *P* = 0.0002; Time spent in chamber by SNI mice Drug x Conditioning interaction F_1,10_ = 118.7, *P* < 0.0001; *n* = 5-6). NPY cKO did not change gabapentin-induced CPP, indicating that the magnitude of ongoing pain experienced by mice following nerve injury is not amplified or attenuated by NPY expression in sensory neurons (**Fig. 3c:** Drug x Injury x Genotype interaction F_1,18_ = 0.311, *P* = 0.58; **Fig. 3d**: Drug x Genotype x Conditioning interaction F_1,10_ = 1.65, *P* = 0.23).

**Figure 3.**
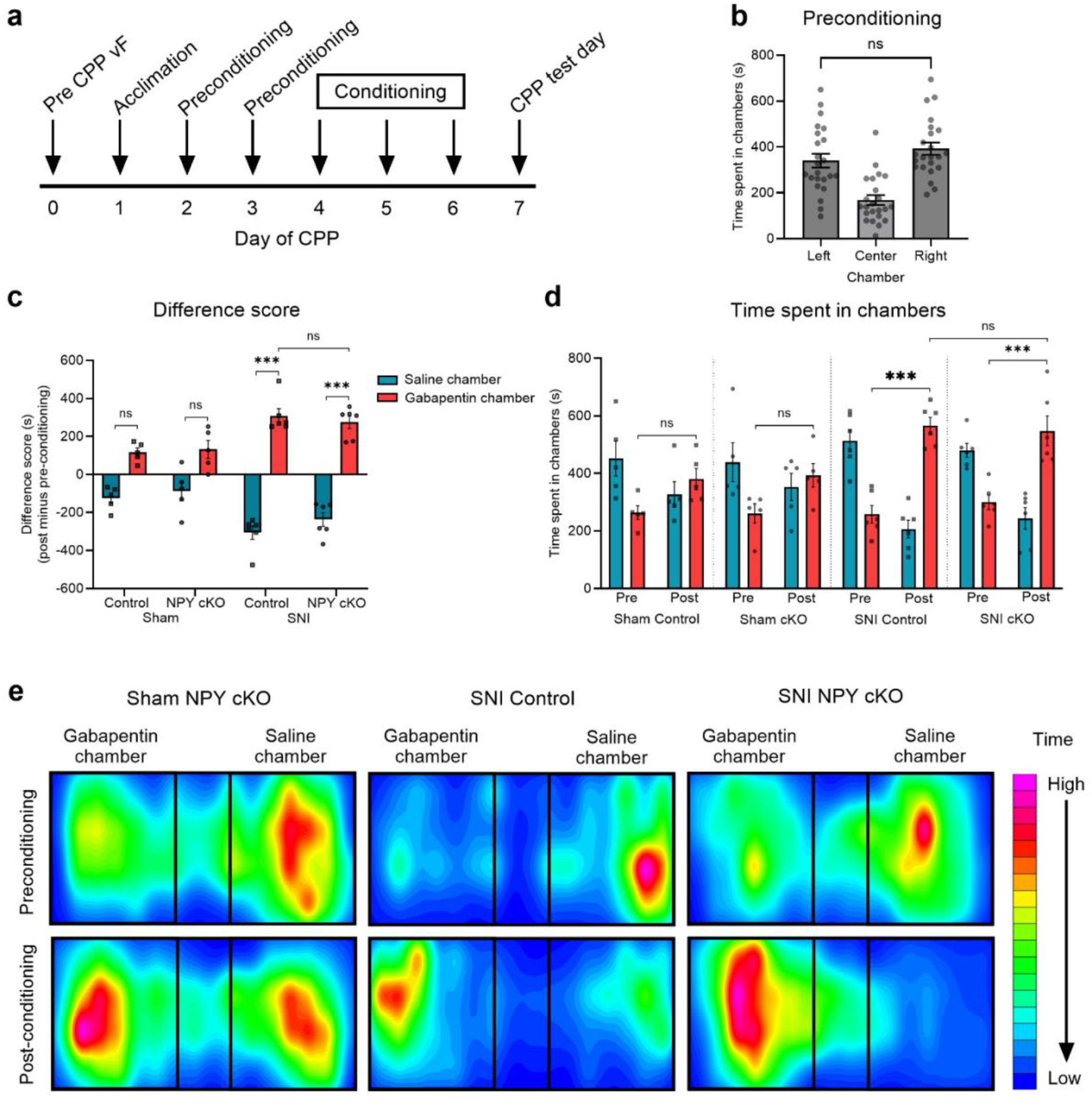
Sensory neuron-specific NPY cKO does not change gabapentin-induced conditioned place preference after SNI. **(a)** Experimental timeline. **(b)** Time spent in chambers during preconditioning (D2 to D3). **(c**,**d)** No difference in gabapentin (100 mg/kg)-induced CPP following SNI was observed between *Pirt*-NPY (NPY cKO) mice and their genetic controls (3-way RM ANOVA with Tukey’s post-tests; *\*\*\* P* < 0.001; *n* = 5-6). **(e)** Representative heat maps demonstrating relative time spent in each chamber during preconditioning and post-conditioning test days, comparing control and *Pirt*-NPY (NPY cKO) mice after sham surgery or SNI surgery. Our biased design explains the greater amount of time spent in the saline chamber during preconditioning. Data represents mean ± SEM.

### Conditional deletion of NPY from primary afferent neurons does not change NPY-mediated suppression of latent pain sensitization

Peripheral nerve injury-induced hypersensitivity resolves into a state of latent pain sensitization due to a compensatory tonic inhibition of nociceptive transmission effected by spinal Y1 and Y2 receptor signaling^11^. To test whether PAN-derived NPY contributes to the endogenous inhibition of latent pain sensitization, we intrathecally administered the NPY Y1R-specific antagonist BIBO3304 (5 µg/5 µL) following the resolution of tSNI-induced hypersensitivity (7 weeks after surgery) in *Pirt*-NPY and control mice. As illustrated in **Fig. 4**, BIBO reinstated mechanical hypersensitivity in tSNI mice but not sham-operated controls (3-way RM ANOVA with Tukey’s post-tests; Time x Drug interaction F_5,110_ = 7.98, *P* < 0.001; *n* = 6-7). NPY cKO did not change reinstatement of hypersensitivity (Time x Drug x Genotype interaction F_5,110_ = 0.83 *P* = 0.53), indicating that sensory neuron-derived NPY does not contribute to tonic inhibition of LS.

**Figure 4.**
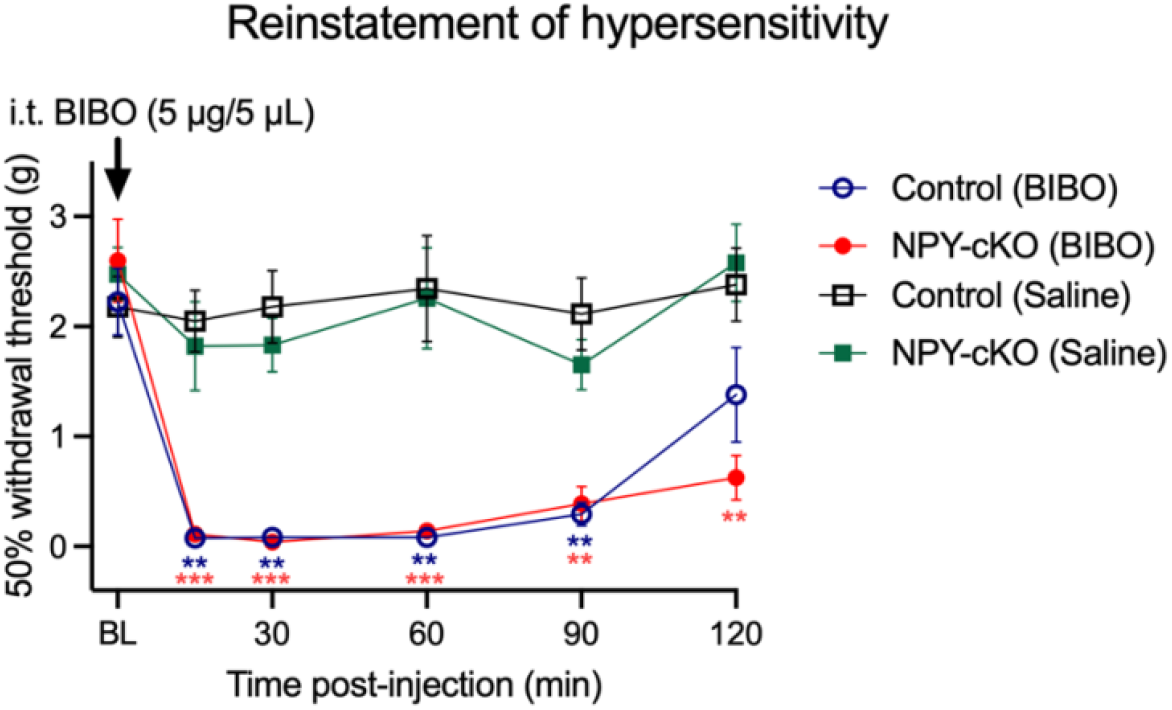
Sensory neuron-specific NPY cKO does not change tonic inhibition of latent pain sensitization by NPY. Mechanical thresholds at the ipsilateral hindpaw following intrathecal injection of the Y1R antagonist BIBO3304 (5 μg/5 μl) or vehicle (saline, 5 μl), in NPY^lox/lox^ controls and NPY cKO mice, 43 days after tSNI surgery. 3-way RM ANOVAs with Tukey’s post-tests; * *P* < 0.05, ** *P* < 0.01, *** *P* < 0.001; *n* = 6-7. Data represents mean ± SEM.

## DISCUSSION

### The *de novo* expression of NPY in sensory neurons is unrelated to neuropathic allodynia

The function of the dramatic upregulation of NPY in myelinated primary afferent neurons and deep dorsal horn following axonal damage^3,36^ has remained an elusive question of great interest for decades. Some studies support a pronociceptive function, notably the finding that microinjection of NPY antiserum or Y1 antagonists into the nucleus gracilis reversed mechanical hypersensitivity following rat sciatic nerve ligation (SNL)^7,8^. By contrast, other studies indicate an antinociceptive function of endogenous NPY. For example, rats overexpressing NPY exhibited reduced behavioral hypersensitivity after spinal nerve ligation^37^, while germline *Npy1r* knockout mice displayed exaggerated allodynia after SNI^10^. We previously showed that conditional *in vivo* NPY^tet/tet^ knockdown of NPY produced a rapid, reversible, and repeatable reinstatement of mechanical and cold hypersensitivity when tested during the remission phase of latent sensitization after tSNI^11^. However, our study used germ-line manipulations leading to NPY knockdown throughout the body, not just sensory neurons. Therefore, we attempted to resolve the function of PAN NPY with a unique genetic approach to selectively delete it from sensory neurons while leaving its expression in CNS neurons intact.

We confirmed that *Pirt*-NPY mice did not exhibit the dramatic up-regulation of NPY in DRG neurons and their terminals in the dorsal horn, while leaving intact NPY expression in the brain and intrinsic spinal cord neurons. Our main finding is that this sensory neuron-specific deletion of NPY did ***not*** reveal any change in neuropathic pain. This was consistent across multiple measure of pain-like behavior, and across multiple peripheral nerve injury models including a model of latent neuropathic pain sensitization. The take-home message of these results is that nerve injury-induced *de novo* expression of NPY in sensory neurons is unrelated to behavioral signs of neuropathic pain or to the endogenous inhibition of latent sensitization. This is not entirely surprising, as our results are consistent with several previous reports. First, Hokfelt and colleagues found that NPY levels in dorsal root ganglia did not correlate with mechanical allodynia in multiple models of peripheral nerve injury (photochemical lesion, partial sciatic nerve ligation, or sciatic nerve constriction)^38^. Second, Colvin and Duggan and colleagues used antibody microprobes to determine that peripheral nerve injury was associated with an increase in the spontaneous release of NPY into deep dorsal that could be further increased by electrical stimulation of large diameter primary afferent neurons^39^, but more importantly found no effect of *in vivo* local anesthetic block injured nerve on this NPY release horn^13^. These results are inconsistent with any contribution of ectopic or nociceptive nerve activity in PANs to spinal NPY release.

### NPY released by DH interneurons, rather than the central terminals of sensory neurons, tonically inhibits spinal neuropathic hypersensitivity

We previously reported that intrathecal injection of NPY Y1R antagonist (BIBO3304) reinstated behavioral signs of inflammatory or neuropathic pain and suggested that endogenous NPY signaling at Y1 receptors in the superficial DH tonically suppresses hypersensitivity^11,40^. Here, we found that after the resolution of tSNI-induced hypersensitivity, intrathecal administration of BIBO3304 reinstated mechanical allodynia to the same extent in *Pirt*-NPY mice as in controls. This result indicates that NPY released by primary afferents does not contribute to the pool of NPY in the spinal cord that tonically inhibits nerve injury-induced hypersensitivity. Notwithstanding a potential contribution of NPY released by descending projections from the brain, we conclude that NPY released by DH interneurons, rather than at the central terminals of damaged afferents, mediates tonic antihyperalgesic signaling following nerve injury. The subset of inhibitory NPY-expressing interneurons^41,42^ that release NPY to tonically inhibit neuropathic pain remains an important unanswered question.

### Further considerations of sensory neuron-derived NPY in neuropathic pain

Our studies contrast with a proposed pronociceptive function of endogenous primary afferent NPY signaling following nerve injury^19^. In addition to the idea noted above that NPY receptor signaling in the nucleus gracilis contributes to mechanical hypersensitivity after SNL^7,8^, injection of the Y2R antagonist BIIE0246 (though not Y1R antagonists) into the DRG blocked the development of SNL-induced mechanical hypersensitivity^9^. As NPY receptors are primarily G_i_-coupled, hypersensitivity might result from disinhibition of intracellular signaling on pronociceptive neurons. For example, studies in dissociated DRG neurons suggest that NPY suppresses N-type Ca^2-^ current via Y1 and Y2 receptors, resulting in attenuation of Ca^2+^-sensitive K^+^ conductance and thus increases in cell excitability. This might explain how nerve injury increases the excitatory action of Y2 agonists^43^. Another putative mechanism is nerve injury leads to NPY-mediated excitation of DRG neurons by inducing a switch in G-protein coupling, as has been observed in µ opioid receptors ^44-46^ and proposed to occur in Y2 receptors ^47,48^. NPY-mediated excitation of DRG neurons may allow NPY to contribute to cross-excitation, a phenomenon in which DRG neurons excite each other via intraganglionic chemical release ^49,50^, thereby increasing nociceptive signaling.

We offer several possible explanations for the discrepancy between previous studies that indicated a pronociceptive contribution of sensory neuron-derived NPY and the current findings of a null contribution. First, nerve injury could activate two NPYergic circuits that negate each other: antinociceptive actions at dorsal horn interneurons that oppose pronociception actions at peripheral terminals, DRG cell bodies, and dorsal column nuclei ^8,9,11^. Second, conditional NPY knockdown may have led to the rapid development of compensatory changes during knockdown. For example, depletion of NPY could be associated with an up-regulation of NPY receptors^51^ or other pain inhibitory circuits that mask the pronociceptive actions of NPY. Third, we note a key species difference: support for pro-nociceptive functions of endogenous NPY signaling following nerve injury were conducted in rats ^7-9^, whilst our study was conducted in mice. Indeed, a lower proportion of DRG neurons express NPY receptors in mice than in rats ^16,20,52-54^, perhaps minimizing the contribution of DRG NPY to neuropathic allodynia in mice. Conversely, an increased proportion of inhibitory DH interneurons express NPY in mice than in rats^41,42^, and thus might enhance the contribution of DH NPY to neuropathic allodynia in mice.

### The *de novo* expression of NPY in sensory neurons does not change ongoing neuropathic pain

The current results indicate that specific deletion of NPY from sensory neurons did not amplify or attenuate gabapentin-induced CPP in nerve-injured mice. These findings are consistent with previous work in which NPY antiserum injected into the nucleus gracilis failed to induce CPP in nerve injured rats ^35^. Furthermore, injection of NPY into the nucleus gracilis failed to produce conditioned place aversion. We conclude that NPY expression in sensory neurons does not contribute to ongoing / spontaneous neuropathic pain.

### The biological function of nerve injury-induced *de novo* expression of NPY in sensory neurons

If not to regulate pain, then what is the biological function of the dramatic up-regulation of NPY in sensory neurons following peripheral nerve injury? We speculate that this NPY contributes to axonal repair and reinnervation. Indeed, NPY promotes neurogenesis in several regions of the CNS, including the dentate gyrus of the hippocampus and the subventricular zone ^55^, as well as sensory neurons within olfactory epithelium ^56^. Interesting results were obtained from cultured DRG neurons obtained from rats with sciatic nerve transection after intrathecal drug injection: first, an NPY antagonist attenuated nerve injury-induced increase in neurite outgrowth; second, NPY increased the percentage of DRG neurons with neurites when co-cultured with spinal cord explants^57^. Finally, Hokfelt and colleagues reported that NPY increased growth rate and regulated axon guidance in embryonic DRG neuron cultures^58^. These studies provide premise for future studies that might use *Pirt*-NPY mice to evaluate the contribution of NPY to axonal repair and reinnervation following peripheral nerve injury.

## AUTHOR CONTRIBUTIONS

*Conceptualization:* A.H.C. and B.K.T.; *Experimental Investigation:* A.H.C. and N.S.H.; *Writing:* A.H.C., A.N., N.S.H. and B.K.T.; *Statistical analysis:* A.H.C.; *Visualization:* A.H.C., A.N. and N.S.H.; *Supervision:* B.K.T.; *Funding acquisition:* B.K.T.

## ACKNOWLEDGEMENTS

R01DA37621, R01NS45954, R01NS62306, and The Raymond and Elizabeth Bloch Educational and Charitable Foundation to B.K.T. We thank Xinzhong Dong for providing us with *Pirt*-Cre mouse breeders.

## Notes

***Conflict of interest:*** The authors declare no competing financial interests

### Competing Interest Statement

The authors have declared no competing interest.

